# Mathematically modelling the population dynamics of CRISPR gene drive systems in the pine pest *Sirex noctilio*

**DOI:** 10.1101/2025.04.12.648502

**Authors:** H. Strydom, R. Ouifki, M. Chapwanya, B. Slippers

## Abstract

*Sirex noctilio* is an invasive pest of pine that has caused significant economic damage in South Africa and many other Southern Hemisphere countries. Current management tools are not efficient in all cases and consequently there is a need for more efficient and targeted control measures. An emerging tool for pest management is the use of gene editing and associated gene drive systems. In this study, we aim to investigate the use of CRISPR-Cas gene drive systems in the management of *S. noctilio* in South Africa. As a first step, we developed a model for the population dynamics of *S. noctilio*, using historical national population monitoring data and incorporating the influence of two main biological control agents of the pest. We then modelled the influence of two different CRISPR-Cas systems on the population dynamics of *S. noctilio* namely, a baseline CRISPR model and Complementary Sex Determination CRISPR (CSD) model. Each model is used to simulate a male and female only introduction strategy to estimate the effectiveness of different methods of introducing the gene drive system. The model calibration was achieved by optimizing the model fit to existing data using the least squares technique. Results suggest that both CRISPR gene drive systems would be effective at controlling the population growth of *S. noctilio* at high levels of introduction, but overall population control would be hindered by practical limitations. Although only two CRISPR models were explored, the underlying population model serves as a framework for further studies into the population dynamics of *Sirex noctilio*, as well as many other CRISPR-Cas gene drive systems.

## Introduction

An emerging technology for the management of pest insects is the use of gene editing and gene drives. Gene based management options have long been used as part of the Sterile Insect Technique (SIT), which uses irradiation to cause DNA mutations that sterilize males (Dyck et al., 2021). Such sterile males can then be released to compete with fertile males. A potentially revolutionary new technology, however, is the emergence of CRISPR-Cas9 gene editing and associated gene drive systems (Singh et al., 2022a,b). Clustered Regularly Interspaces Short Palindromic Repeats or CRISPR for short is a simple immune system found in bacteria and some archaea (Jansen et al., 2002; Makarova et al., 2002). This system has in recent years been modified to serve as a gene editing technology. Applications include gene knockout and knock-in, gene editing and modification, and in recent years as a very effective gene drive systems (McFarlane et al., 2018). CRISPR gene drives function by using the cells’ natural DNA repair systems to copy itself into apposing chromosomes (Klipp et al., 2016). This results in super-mendelian inheritance of the gene drive making it useful for population modification or suppression.

Many insect species have been modified using CRISPR technology and many studies are looking into possible applications in population control of insects using this technology (Singh et al., 2022a; D. Sun et al., 2017; Yan et al., 2022). One of the major focus areas of population control using CRISPR technology at present is on the management of insect and disease vectoring pests (Yan et al., 2022). Research has predominantly concentrated on species that transmit human diseases due to their significant public health impact (Dong & Dimopoulos, 2023; Edwards, 2023; Nuss et al., 2023; Papathanos et al., 2024). However there has also been a focus on studying plant pests, reflecting the importance of protecting agriculture from these harmful species (Komal et al., 2023; Moon et al., 2022; Singh et al., 2022a).

Ensuring that a CRISPR gene drive is safe and effective requires a significant amount of research. This is because gene drives must be tailored to the specific organism being targeted and special care must be taken to prevent the spread of the gene drive outside the intended population (Esvelt & Gemmell, 2017; Marshall, 2009; Noble et al., 2018). For example, using mathematical modeling, Faber, McFarlane, et al. (2021) demonstrates how a combination of different CRISPR systems can be used to control an invasive squirrel population. This system reduces the rate of resistance development and helps localize the spread to a single population. This showcases how mathematical models can help alleviate some of these issues by allowing researchers to simulate many different possible scenarios, developing hypotheses, and directing research (Klipp et al., 2016).

*Sirex noctilio* is an invasive woodwasp that has been introduced into many countries around the world (Slippers, et al., 2012). This woodwasp was first introduced into New Zealand, followed by Australia and it eventually emerged in South America, South Africa, North America and most recently in China (Slippers, de Groot, et al., 2012; X. Sun, Tao, et al., 2020). This insect originates from Eurasia, where it is considered a minor pest. *Sirex noctilio*, together with its fungal symbiont, *Amylostereum areolatum*, has, however, become a major pest of commercial pine tree plantations in many of the regions where it has been introduced.

Management of *S. noctilio* has proven difficult and methods of controlling the spread and reducing the economic impact of this pest has not always been efficient (Corley et al., 2019; Hurley et al., 2012). This is partly because of the cryptic life cycle of *S. noctilio*. For most of its life, the larvae of the wasp is protected inside the trunk of trees it has infested. Adults only emerge for a few weeks during the end of the life-cycle when mating and reproduction takes place. In many countries suppression has been achieved by use of biological control agents, and in particular, the nematode *Deladenus siricidicola* (Slippers et al., 2012) and the parasitic wasp *Ibalia leucospoides* (Cameron, 2012). *Deladenus siricidicola* is a parasitic nematode that infects and invades the reproductive system of *S. noctilio* rendering females sterile. *Ibalia leucospoides* is a parasitic wasp that targets the eggs or early instar larvae of *S. noctilio*. Although these two biocontrol agents have been successful in many cases, they could not prevent outbreaks in all cases. There are also concerns about the sustainability and long-term effectiveness due to environmental variability, natural resistance or evolution of resistance. More research is thus required into alternative methods of control.

In this study, we explore the use of mathematical modeling to test hypotheses for the use of CRISPR technology for the management of *S. noctilio* populations. We firstly develop a model consisting of a system of ordinary differential equations (ODEs) that describe the population of *S. noctilio* populations in South Africa, based on historical monitoring data and published research. The model also includes the influence of *D. siricidicola* and *I. leucospoides*. Then, we incorporate into the model the effects of CRISPR gene drive systems on these populations using the baseline population dynamics model. We explore two approaches, namely a generic approach to population suppression, as well as an approach that targets the Hymenoptera/*S. noctilio* specific Complementary Sex Determination reproductive biology system.

## Materials and Methods

### Data description

Two datasets were available for *S. noctilio* sourced from nine distinct regions spanning South Africa. The data was gathered as a component of a national initiative aimed at monitoring the distribution of *S. noctilio* and evaluating the effectiveness of introduced biological control agents, overseen by the National Sirex Steering Committee (Philip Croft, personal communication).

Data collection for this dataset involved the felling of *S. noctilio* infested trees, cutting them into 80 cm billets and placing the collected logs into cages. Any adult *S. noctilio* or *I. leucospoides* that emerged were collected. *Sirex noctilio* adults were dissected to determine infection by *D. siricidicola*. The total amount of females, females infected with *D. siricidicola*, males, males infected and adult *I. leucospoides* were recorded per log.

In addition, countrywide infection surveys were conducted in known *S. noctilio*-infested plantations, during which the count of living (uninfected), dying (infected), and dead (from the previous season) trees was recorded.

Data was available for nine regions of South Africa from 2007 to 2017: Western Cape, Eastern Cape, Southern Cape, KwaZulu-Natal North, South and Midlands, Limpopo, Mpumalanga North, and Mpumalanga South. Due to the nature of how the data was collected few regions had data available for consecutive years. We chose to use the KwaZulu-Natal Midlands as it had the longest continuous stretch of data from 2012 to 2017 with 1393 trees surveyed. The data from this region was used to calculate model parameters and population sizes for each year. The remaining model parameters were estimated through fitting our proposed model to the available data using a least square fitting process.

The population size per tree was calculated, and the data integrated with survey results to provide estimates for the overall population size of both *S. noctilio* and *I. leucospoides*, as well as an estimation of the infection rate for *D. siricidicola*. To facilitate subsequent calculations, the population size was scaled down to a representation based on 1000 trees. This was done to prevent overly long computational times. Trees infected with *S. noctilio* die, preventing reinfection and decreasing the number of trees available. This was considered when calculating the population sizes.

### Population model

We start by developing a model for the population dynamics of *S. noctilio*, that accounts for the effects of two main biological control agents of the pest. The life cycle of *S. noctilio* includes non-adult stages (eggs, larvae, and pupae) and adult stages. Non-adult stages are predominantly confined within the host tree, we combine them in our model into a unified “juvenile” category. Adults only emerge for a few weeks to reproduce before dying. Infection of *S. noctilio* with *D. siricidicola* leads to the sterility of females and transforms them into carriers of the entomopathogenic nematode (Bedding, 1972). Infected and uninfected males are biologically indistinguishable but are treated as separate groups in our model for completeness. Subsequently, we divide the population of *S. noctilio* into eight distinct groups based on life stage, gender, and infection status. These categories encompass males and females, adults and juveniles, and individuals infected with *D. siricidicola* or uninfected (normal).

A comprehensive list of all eight variables and their descriptions are given in Table 1.

**Table 1.** Population model variables for *S.noctilio*.

| Variable | Description |
| --- | --- |
| $A_{f_N}$ | Adult, female, uninfected |
| $A_{fD}$ | Adult, female, infected |
| $J_{fN}$ | Juvenile, female, uninfected |
| $J_{fD}$ | Juvenile, female, infected |
| $A_{mN}$ | Adult, male, uninfected |
| $A_{mD}$ | Adult, male, infected |
| $J_{mN}$ | Juvenile, male, uninfected |
| $J_{mD}$ | Juvenile, male, infected |

To describe the population dynamics of *S. noctilio* we develop a deterministic model consisting of eight differential equations that describe the interaction dynamics between these eight classes (Equation 2). The corresponding model diagram is presented in Fig. 2. In our model formulation we account for the following consideration:

- *S. noctilio* population:
  ∘ *S. noctilio* is an arrhenotokous or haplodiploid species, wherein males are haploid and originate from unfertilized eggs, while females are diploid and result from sexual reproduction (Slippers, de Groot, et al., 2012).
  ∘ The egg-laying rate of the population is encapsulated in the parameter ε.
  ∘ It is worth noting that diploid males do occur naturally but constitute only a small fraction of the population, and they are sterile, so they are omitted from our model (Queffelec et al., 2019).
  ∘ We employ two time-dependent functions to govern the life cycle of *Sirex noctilio*. The emergence function, denoted as *w*(*t*), oversees the transition from the juvenile stage to adulthood, while the reproduction function, represented as *σ*(*t*), represents the reproduction rate of the adult population (Fig. 1). Adults die after reproducing.
  ∘ It is important to note that these two functions occur in rapid succession without any overlap. This aims to prevent the occurrence of a feedback loop in which newly produced juveniles would be capable of reproduction.
  ∘ Towards the end of the year, during the emergence period, *w*(*t*) exhibits a notable increase, approaching a value of 1. This surge in *w*(*t*) facilitates the transition of juveniles into the adult stage, with a propotion *χ* successfully transitioning to adulthood. Subsequently, *w*(*t*) returns to its baseline value of 0. *σ*(*t*) follows similar trends with a slight delay precluding any overlap with the emergence function *w*(*t*).
  ∘ The interplay between sexual and asexual reproduction is captured by the function *ϕ*(*p*), proposed by Queffelec et al., 2019

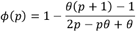

where represents the adult male-to-female ratio within the population given by

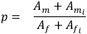
  ∘ For most of the year, both *σ*(*t*) and *w*(*t*) remain equal to 0, corresponding to the period where *S. noctilio* is in its juvenile developmental phase.
  ∘ Moreover, the model accounts for various sources of mortality, encompassing juvenile mortality (*µ*_*j*_), adult male mortality 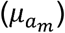, and adult female mortality 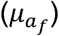. Given that adults die shortly after reproducing, we opt to model the temporal mortality rate as *μ*_*c*_*σ*(*t*), where *c* represents *j*,*a*_*m*_ or *a*_*f*_.
  ∘ The model does not incorporate migration as a contributing factor, primarily due to the unavailability of relevant data.
- Biological control:
  ∘ The model includes two biological control agents that are applied in South Africa: the infection rate of *S. noctilio* by *D. siricidicola* denoted by *α*, and parasitism by *I. leucospoides* denoted by *γ*.
  ∘ A logistical function was developed to estimate *α*. Our analysis revealed that the maximum infection rate, 48%, is attained when the proportion of infected females within the *S. noctilio* population reaches 16%. Consequently, this logistic function exhibits an asymptote at *α* = 0.48 and an inflection point at *a* = 0.08. To ensure a minimal rate of infection in the absence of infected females, the function was left-shifted to ensure *α*(0) is approximately 0. Thus, the proportion of infected individuals can be expressed as:

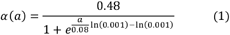

where *a* is the proportion of infected adult females to the total population size given by

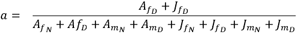
  ∘ We also observed a one-to-one relationship between the presence of *I. leucospoides* individuals and *S. noctilio* mortality. We assumed a constant parasitism rate (*γ*), dependent solely on the total population size of *S. noctilio*.
  ∘ Our analysis of the available data indicated negligible variations in infection and predation rates between male and female *S. noctilio* individuals.

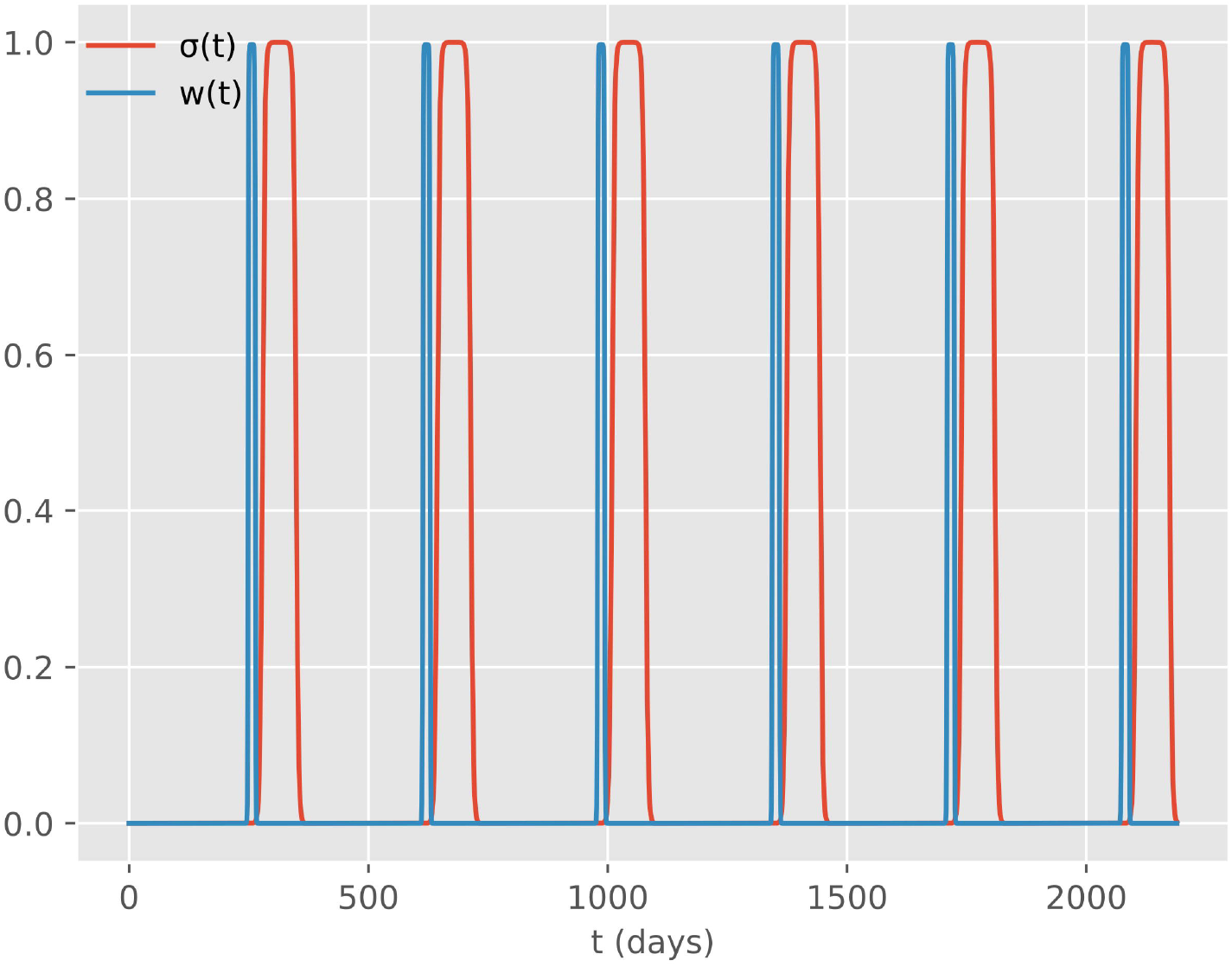

The model diagram (Fig 2) shows all the variables, parameters, and interactions in our model. The diagram is divided into three panels: Panel (A) represents the reproduction phase where fertile female adults reproduce several offspring (*ϵ*) either sexually (*ϕ*(*p*)), producing male and female offspring, or asexually (1 − *ϕ*(*p*)) producing only male offspring. The process of reproducing is controlled by the reproduction function (*σ*(*t*)) and takes place over two weeks after emergence has occurred. Panel (B) illustrates the infection phase where a proportion of the juvenile population is infected by *D. siricidicola* (*α*(*a*)). Infected females are sterile while infected males are not. (C) depicts the final emergence phase of the juvenile population transition into adults. This process is controlled by the emergence function (*w*(*t*)) and takes place over two days right before reproduction occurs.

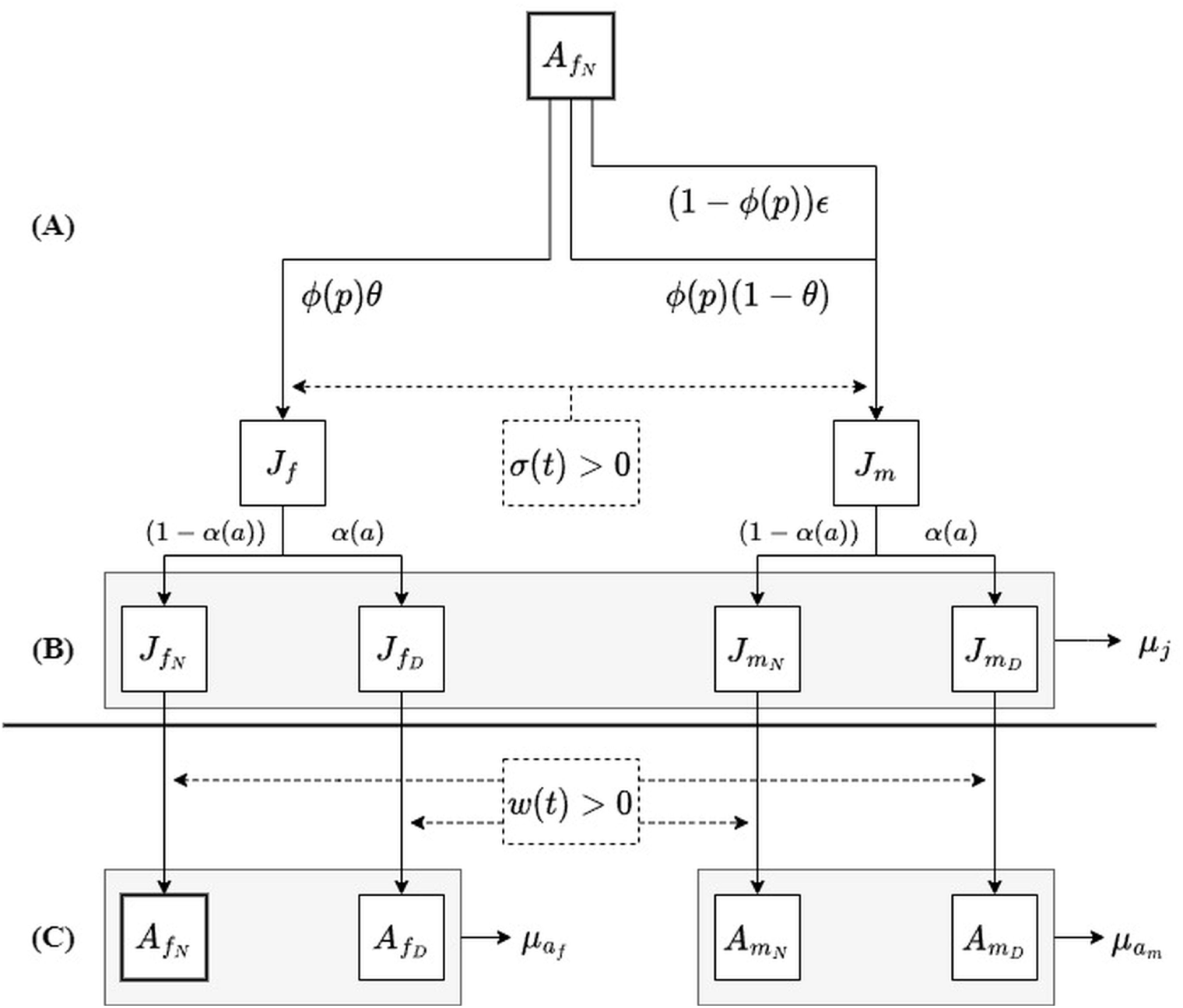

The final model equations are summarised as follows (Fig 2):

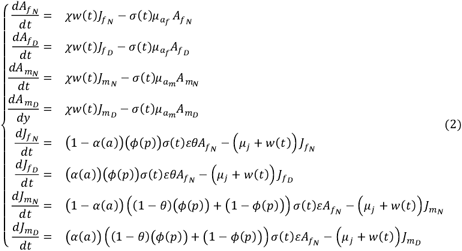

### Parameter optimization

Model parameters were optimized for summer rainfall regions of South Africa, but the underlying mathematical model is generic and is applicable to any population.

The dataset encompasses records spanning from 2012 to 2017. Various parameters were derived from this dataset, including the sex ratio (θ), the prevalence of *D. siricidicola* infection among adult *S. noctilio* individuals, the predation rate of *I. leucospoides* (*γ*), and the initial population sizes for each variable. Mortality rates were estimated based on the average number of days an individual is expected to survive each year. Specifically, the juvenile stage persists for 351 days, whereas adult males have a lifespan of 12 days, and adult females a mere 5 days (Ryan & Hurley, 2012). Egg-laying capacity is intricately linked to female body size and can range from 30 to 450 eggs laid over the course of a female’s lifetime (Haavik et al., 2017; Madden, 1974). Notably, the estimates for egg-laying capacity and initial population sizes underwent refinement as part of the model fitting process.

The model calibration was conducted utilizing the least squares (leastsq) optimization method, implemented with the lmfit package version 1.2.2 in Python, version 3.9.16. The optimization process focused on determining the values of the egg-laying rate (ε) and the initial conditions for four key variables 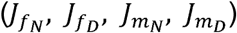.

To validate the model’s performance, its predictions of population size were compared to the estimated population sizes obtained from empirical data. The goal of the calibration process was to minimize the differences between the model’s predictions and the observed data. During this fitting procedure, 95% confidence intervals were considered to ensure the robustness of the model’s parameter estimates.

The available dataset contained information that allowed the estimation of peak population sizes on an annual basis across a span of six years, encompassing four distinct variables.

### CRISPR models

The integration of two CRISPR gene drive systems into our models follows a deterministic Susceptible-Infected-Recovered (SIR) framework, building upon the existing population model without altering any pre-established parameter values. The key addition is a singular parameter, homing efficiency (δ), which quantifies the ability of the CRISPR construct to insert itself into a susceptible allele. It is assumed that alleles failing to undergo transformation inherently become resistant.

The individuals in the population are categorized by their genotype, with three allelic states: Susceptible (s), representing wild-type individuals; Infected (i), embodying individuals carrying the theoretical CRISPR gene drive construct; and Resistant (r), indicative of individuals resulting from non-homologous end joining events occurring in infected individuals. This allelic diversity results in six female genotypes (*ss, si, sr, ii, ir, and rr*) and three male genotypes (*s, i, and r*).

### Model generalisation

Generalising the original population model equations (Equation 2), we can describe any individual in a population using four equations.

A diploid (female) individual can be describe using:

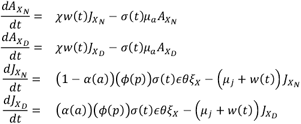

A haploid (male) individual can be described using:

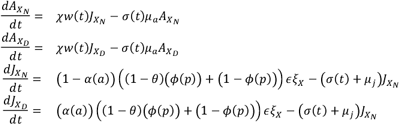

In both cases, *X* denotes the individual’s genotype and *ξ*_*X*_ the proportion of offspring in the population expected to have the genotype *X*. *ξ*_*X*_ changes for each genotype, based on the CRISPR gene drive mechanism being modeled.

The frequency of any male in the population can be expressed as:

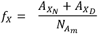

Where *X* is the genotype and 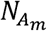 is the total, adult male population.

### Offspring frequencies

Offspring genotype frequencies were determined by summing the probabilities of all mating combinations that could produce each specific genotype. For instance, in a wild population with three alleles (*X, Y and Z*) segregating according to Mendelian inheritance, the frequency of producing a female offspring with genotype *XX*(*ξ*_*XX*_) depends on mating between a female carrying at least one *X* allele (*A*_*XX*_, *A*_*XY*,_ *A*_*XZ*_) and a male carrying an *X* allele (*f*_*X*_).

The offspring frequency for *XX*(*ξ*_*XX*_) is therefore the sum of the probabilities of the following mating scenarios:

1. An *A*_*XX*_ female and a *f*_*X*_ male produce one possible *XX* offspring:

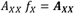
2. An adult female (*A*_*XY*_) mates with a male (*f*_*X*_) resulting in two possible offspring, one of which is *XX*:

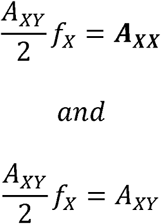
3. An adult female (*A*_*XZ*_) mates with a male (*f*_*X*_) resulting in two possible offspring, one of which is *XX*:

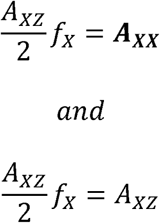

Thus *ξ*_*XX*_ can be calculated as:

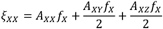

By summing these probabilities for each parental genotype pair, we obtain the overall offspring frequencies in the population.

When introducing homing efficiency, alleles can transform other alleles at a rate ***δ***. This homing modifies allele frequencies such that combinations involving the homing allele increase by a factor of (**1 + *δ*)**, while corresponding alleles without homing decrease by (**1 − *δ*)**, thereby altering the genotype frequencies.

### Baseline CRISPR model

The baseline model assumes a CRISPR construct where a single allele confers no negative effect, while diploid individuals are considered sterile. Resistant individuals do not contain the deleterious construct and suffer no decrease in fitness.

Offspring frequencies for the model were calculated as:

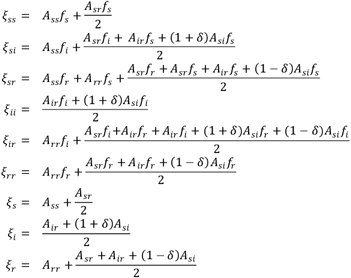

### Complementary sex determination model

The CSD model assumes a CRISPR construct that flanks the sex determination locus of *S. noctilio. Haploid C*RISPR individuals are unaffected and are phenotypically indistinguishable from wild-type individuals. Diploid CRISPR individuals develop as sterile males due to having two of the same CSD loci. Resistance alleles are alleles that do not contain a sex determination locus, individuals carrying a resistant allele are assumed to be sterile.

Offspring frequencies for the model were calculated as:

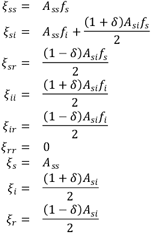

## Results

### Population model

Adult population sizes were estimated from the available KwaZulu-Natal Midlands data for each consecutive year (2012 to 2017), for each of the estimates a 95% confidence interval were also calculated. The estimate for infection with *D. siricidicola* at population saturation was calculated at 48 % of newly emerged adult *S. noctilio*. Predation rate by *I. leucospoides* was calculated at 33 % of the juvenile population and population sex ratio was calculated at 34 % female. Mortality rates were derived from biological knowledge, with juvenile mortality per day (*µ*_*j*_) = 0.00284, adult female mortality (*µ*_*f*_) = 0.2 and adult male mortality (*µ*_*m*_) = 0.083.

Wellness of fit was calculated using a chi-square test. A chi-square value of 26.79 with 19 degrees of freedom were obtained, with a resulting p value of 0.1095. The model predicts a decrease in total population size across time, with initial values for the juvenile population at 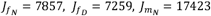 *and* 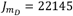. Egg laying capacity (*ϵ*) was predicted at 4.56 eggs per day.

Visual representations of the model fit, containing the model predictions, data estimates and confidence intervals can be found in Fig 3. All parameters, their final estimates and descriptions can be found in Table 3. Functions and their descriptions can be found in Table 4.

**Table 2.** Adult Population Estimates.

| YEAR | POPULATION ESTIMATE |  |  |  |
| --- | --- | --- | --- | --- |
| | $A_{f_N}$ | $A_{f_D}$ | $A_{m_N}$ | $A_{m_D}$ |
| 2012 | 2755 ± 809 | 2520 ± 828 | 5635 ± 1358 | 7000 ± 2157 |
| 2013 | 2934 ± 728 | 2031 ± 502 | 9103 ± 2302 | 6868 ± 1854 |
| 2014 | 2921 ± 1030 | 2326 ± 1206 | 5126 ± 1490 | 3921 ± 1997 |
| 2015 | 3427 ± 917 | 1951 ± 763 | 5489 ± 1160 | 2363 ± 756 |
| 2016 | 3440 ± 1012 | 1839 ± 633 | 6398 ± 1658 | 6431 ± 2255 |
| 2017 | 1899 ± 556 | 3803 ± 936 | 4132 ± 1163 | 9317 ± 2517 |

**Table 3.** Population model parameter estimates.

| Paramater | Estimate | Description |
| --- | --- | --- |
| $\theta$ | 0.34 | Proportion of offspring that develop into female adults. |
| $\gamma$ | 0.33 | Proportion of juvenile individuals predated by *I. leucospoides*. |
| $\epsilon$ | 4.56 | Total amount of eggs laid per day. |
| $\mu_j$ | 0.00284 | Mortality rate of juvenile individuals per day. |
| $\mu_{af}$ | 0.20 | Mortality rate of adult female individuals per day. |
| $\mu_{am}$ | 0.083 | Mortality rate of adult male individuals per day. |

**Table 4.**
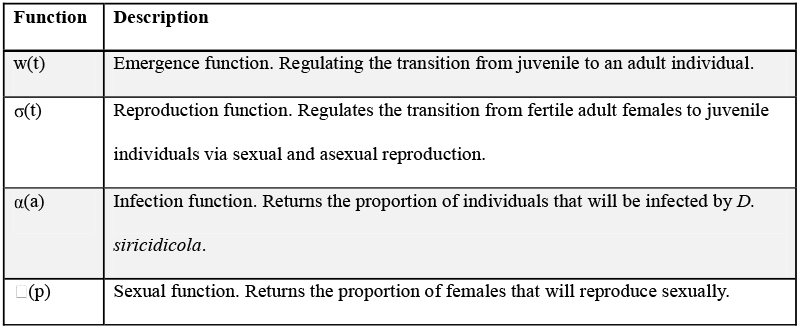
Population model functions.

| Function | Description |
| --- | --- |
| $w(t)$ | Emergence function. Regulating the transition from juvenile to an adult individual. |
| $\sigma(t)$ | Reproduction function. Regulates the transition from fertile adult females to juvenile individuals via sexual and asexual reproduction. |
| $\alpha(a)$ | Infection function. Returns the proportion of individuals that will be infected by *D. siricidicola*. |
| $\square(p)$ | Sexual function. Returns the proportion of females that will reproduce sexually. |

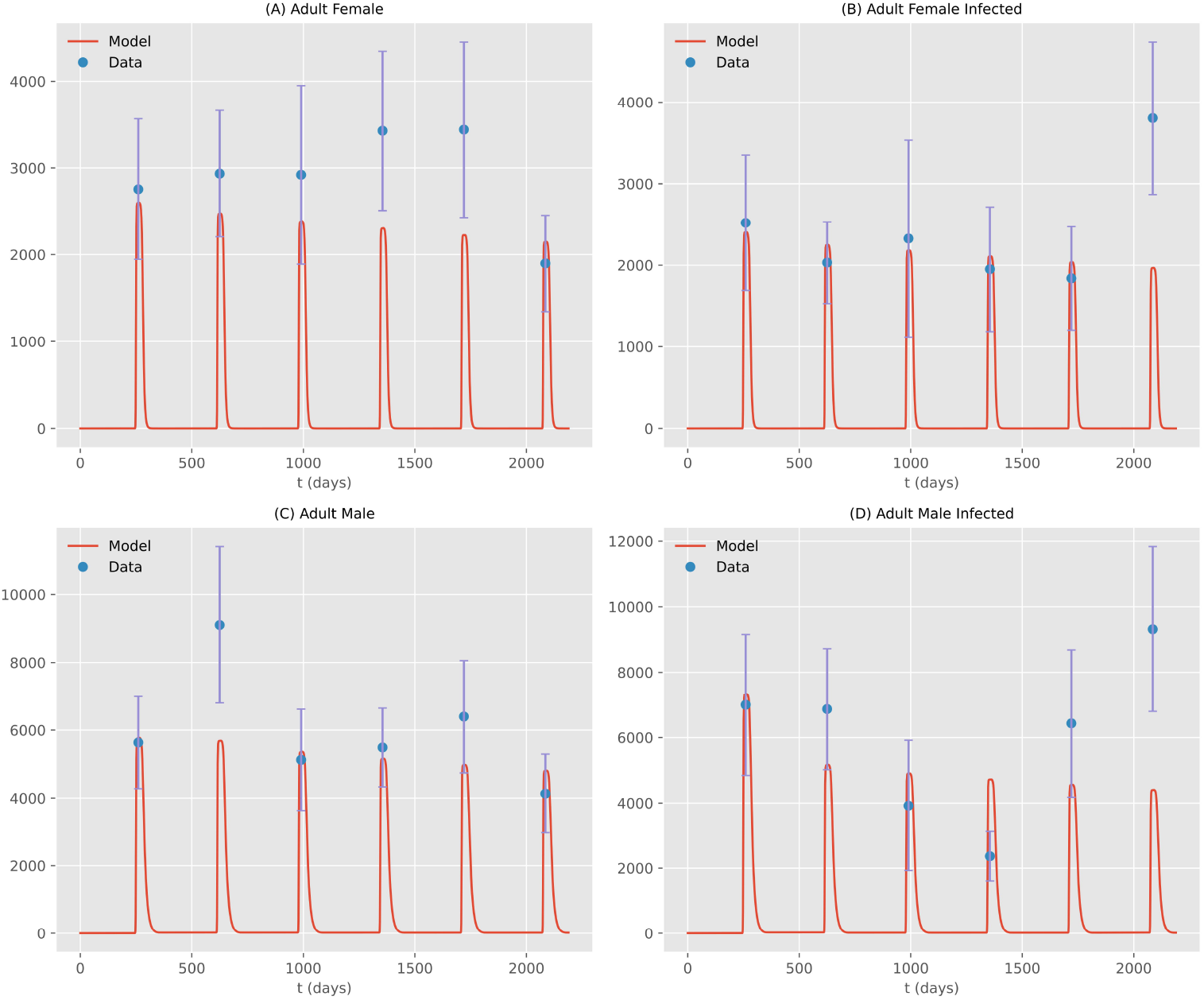

### CRISPR Models

To assess the efficacy of the two CRISPR gene drive strategies, we conducted numerical simulations for each, introducing either only female CRISPR-carrying individuals or only male individuals. The simulations encompassed a range of introduction percentages, spanning from 0% to 100% of the population size in 1% increments. For male introductions, this ranged from 0 to 17,423 individuals, while for females, the range extended from 0 to 7,857 individuals.

Within each population step, the homing efficiency parameter (δ) was systematically varied from 0 to 1 in 0.1 increments. The outcome of each simulation was evaluated by comparing the population size in the final year (N_F_) to the initial population size (N_0_). The resultant change in population size (ΔN) across all permutations was visualized as a heatmap.

The heatmap results for the baseline model are represented in Fig 4. The results for the CSD model are represented in Fig 5.

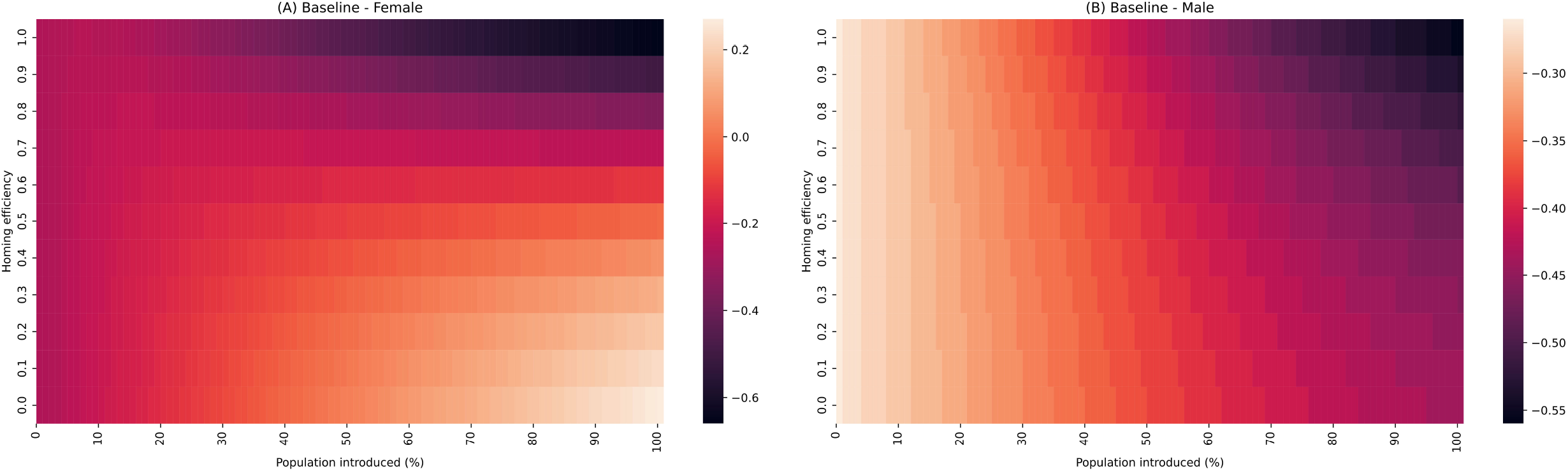

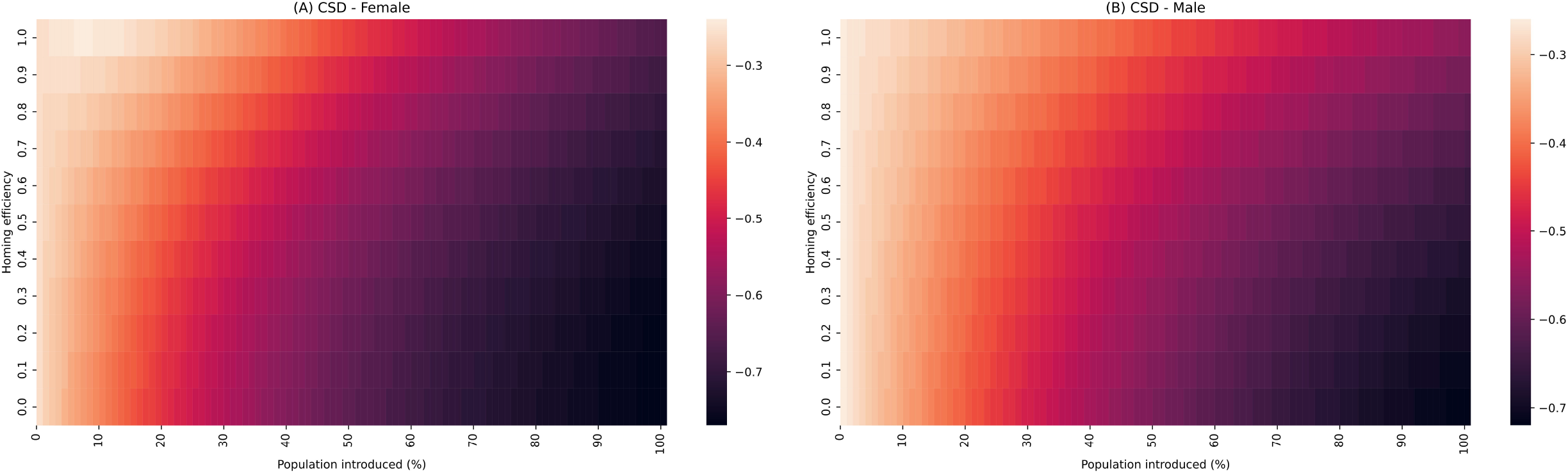

The equation used to determine the change in population size across all permutations was:

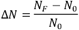

Results for the Baseline CRISPR model show a clear trend in population suppression. The female introduction strategy (Fig 4A) shows a trend where an increase in individuals introduced leads to a decrease in overall population suppression (27% population growth, compared to the baseline 26% population decrease). The combination of increased individuals introduced coupled with higher homing efficiencies does still produce the best results, with total population size decrease of 61%. For the male introduction strategy (Fig 4B) increasing the number of individuals introduced, or increasing the homing efficiency of the drive, increases the total amount of population suppression observed. At the upper limits this correlates to a 66% decrease in population growth across the simulated six years.

Results for the CSD model show similar trends for both the female and male introduction strategy. Increasing the number of individuals introduced improves overall population suppression, but increasing the homing efficiency has a negative effect reducing the population suppression. The female introduction strategy (Fig 5A) does however outperform the male introduction strategy (Fig 5B) with a decrease in population size of 77% at *δ* = 0 and introduction = 100% compared to 72% for the males. At maximum homing efficiency the population suppression drops down to 66% for the females and 56% for the males.

## Discussion

In this study, the first comprehensive population model for *S. noctilio* was developed, enabling dynamic simulations of population growth over time. Our model uniquely incorporates the influence of two prominent biological control agents, *D. siricidicola* and *I. leucospoides*. While models for *S. noctilio* do exist, they predominantly focus on predicting the spatial spread (Gao et al., 2021; Gao & Shi, 2021; Munro et al., 2022; X. Sun, Xu, et al., 2020) or identifying potential susceptible hot spots in plantations (Abdel-Rahman et al., 2014; Fischbein et al., 2019; Ismail et al., 2010; Ismail & Mutanga, 2010; Yemshanov et al., 2009). The model developed here addresses a critical gap by simulating the population dynamics and interactions with key biological control agents. Previous studies have delved into specific aspects such as the spread of *S. noctilio* biological control agents, the emergence of the species, and factors influencing sex determination (Corley et al., 2019; Haavik et al., 2013; Queffelec et al., 2019). This integrated approach, however, expands the scope, offering a holistic understanding of the population dynamics of *S. noctilio* in the context of its biological control agents.

A notable limitation in the study of population dynamics of *S. noctilio* is the scarcity of research on the infection dynamics of *D. siricidicola*. While the biological aspects of the nematode are well-documented (Bedding, 1972; Castillo et al., 2018; Slippers, Hurley, et al., 2012), there is a scarcity of mathematical descriptions of its dynamics. Our study only had access to *S. noctilio* dissection data, indicating whether a given individual was infected or not. This gave indicates the maximum infection rate in the population, but not how that infections rate would change with the population. In our model, we address this gap by assuming a standard logistical curve (Equation 1) as used to model infection in other systems (Dukic et al., 2013; Gupta & Vale, 2017). However, it is important to acknowledge that these assumptions may not hold universally. Changes in parasitism rate, biological control programs and climate will have an influence on the dynamics of *D. siricidicola* infection. More comprehensive data and targeted research on *D. siricidicola* would greatly enhance the accuracy and applicability of this aspect of our model.

The population model serves as the fundamental framework upon which the CRISPR models were developed, providing a foundation for the exploration of various CRISPR constructs and gene drive systems. The lack of data available on CRISPR in *S. noctilio*, makes testing our underlying CRISPR models difficult. We opt to test differing extremes to see if the models perform as expected. To this end we created two contrasting theoretical gene drive systems to demonstrate the versatility of the underlying models. A baseline model based on a haploinsufficient underdominance system (Harvey-Samuel et al., 2017) that allows resistance to develop freely and a complementary sex determination (CSD) model that assumes a gene drive system that targets the sex determination genes of *S. noctilio* where no resistance is allowed to develop. Similar systems have been tested in *Anopheles gambiae* (Kyrou et al., 2018), *Drosophila melanogast* (Chen et al., 2023) and *Drosophila suzikii* (Yadav et al., 2023) and were found to be effective at population suppression.

The development of resistance in CRISPR gene drive system has been known for a while to be one of the main limiting factors to the effective use of gene drive systems (Champer et al., 2018; Price et al., 2020; Unckless et al., 2017). The baseline model performed as expected of a gene drive system with freely developing resistance. Our model demonstrated how lower homing efficiencies, and thus higher rates of resistance development, results in lower rates op population suppression, while higher homing efficiencies resulted in more efficient population suppression.

In the CSD model, we simulate more modern CRISPR gene drive systems, where the development of resistance is actively prevented or penalised by drastically reducing the fitness of resistance carrying individuals. Many similar systems have been suggested, including cleave-and-rescue systems (Faber, McFarlane, et al., 2021; Oberhofer et al., 2019), sex distortion systems like X/Y shredders (Alcalay et al., 2021) and systems that target reproductive genes (Chen et al., 2023; Kyrou et al., 2018; Xu et al., 2020; Yadav et al., 2023).

Our model predictions agree with results found by Faber, McFarlane, et al. (2021), where meaningful population suppression only occurs at high levels of introduction and where individuals show reduced reproductive fitness at higher levels of homing efficiency.

It is important to acknowledge that while our models explore a broad spectrum of introduction percentages and homing efficiencies, practical considerations in field conditions impose significant limitations. When considering a homing efficiency of 90 % (Champer, Yang, et al., 2020) with an introduction range of 1-20 %, both models appear incapable of causing meaningful suppression of the population. This aligns with findings from other researchers (Liu & Champer, 2022) who have demonstrated that even under ideal conditions, many standard gene drive systems, including sex-based (Faber, Meiborg, et al., 2021; Lester et al., 2020) and toxin-antidote drives (Champer, Champer, et al., 2020; Champer, Kim, et al., 2020; Champer, Lee, et al., 2020), struggle or fail to achieve population suppression. Other studies also show that only specific gene drive systems perform well in haplodiploid species (Liu & Champer, 2022). These difficulties highlight the need for mathematical models to inform the development and eventual testing of efficient gene drive system.

While this study specifically investigated the population dynamics of two CRISPR gene drive systems in *S. noctilio* within the context of South Africa, the models established are applicable to any population of *S. noctilio*. The underlying principles and parameters employed in the models are not limited to the South African population but can be adapted for the study of *S. noctilio* populations in different geographical locations. Additionally, the models can serve as a versatile framework for examining the potential outcomes of diverse CRISPR gene drive strategies in *S. noctilio*.

